# Zika virus may not be alone: proteomics associates a bovine-like viral diarrhea virus to microcephaly

**DOI:** 10.1101/062596

**Authors:** Fabio CS Nogueira, Erika Velasquez, Adriana SO Melo, Gilberto B Domont, Akira Sawa

## Abstract

**One Sentence Summary:** Proteomics analysis lead us to suspect the presence of a Bovine-like viral diarrhea virus (BVDV-like) in the brain tissue of fetuses bearing microcephaly during the outbreak in Paraíba State, Brazil, 2015.

**Abstract:** No direct experimental causal evidence confirms that the Zika virus is the sole etiological agent responsible for the development of brain malformations in human fetuses during pregnancy. We used a discovery-driven approach to analyze protein extracts of three Zika positive brains. Shotgun mass spectrometry (MS) proteomics did not identify any Zika protein in all samples. However, MS detected the presence of peptide(s) from the polyprotein of a Bovine-like viral diarrhea virus (BVDV-like) in Zika-positive brains. These results indicate that Zika virus may not be, per se, the only etiological agent responsible for microcephaly and suggests that discovery-driven approaches play an essential role in the screening of fluids or tissues for virus or other etiological agents.

---

To the Zika infection in Bahia, Brazil in 2015 (1) followed a dramatic outbreak of microcephaly in the Northeastern Brazilian States of Paraíba, Pernambuco and Bahia (2). In January 2016, Melo suggested a correlation between Zika infection and fetal microcephaly during a two case study of women living in Paraiba State who complained of symptoms of Zika virus infection during pregnancy; real time PCR confirmed the presence of Zika virus in children brain tissue and amniotic fluid (3). Other groups associated the presence of Zika virus to microcephaly (4) and reviewed the evidence for causality by several criteria (5). However, reports linking high incidence of human microcephaly to the presence of Zika virus do not disclose any experimental data that provide a direct link between microcephaly and Zika virus during fetal brain development in spite of reducing viability and growth of neurospheres and brain organoids (6). Here, we expand these findings using discovery-driven shotgun proteomics based on high resolution and accuracy mass spectrometry analysis to show the detection of a BVDV-like virus in tissue extracts of Zika positive brains.

Microcephaly is associated with genetic disorders. Among others, chromosomal trisomy (Edward’s), Aicardi-Goutieres´ and Rett´s syndromes and X-chromosomal microcephaly and trans-placental infections by viruses or bacteria, including rubella, cytomegalovirus, herpes simplex, varicella zoster virus, HIV, arboviruses and maternal malnutrition or drug use and chemical intoxication of pregnant mothers (3–5, 7).Reports disclosed the incidence of Central Nervous System (CNS) malformations in fetuses and neonates after the Zika virus outbreak in French Polynesia; however, incidence of microcephaly associated with the outbreak was not documented (6). In 2016 was published the complete genome of Zika virus sampled from the amniotic fluid of fetuses with microcephaly living in Brazil´s northeast (8).

Some questions are still unanswered as the association between Zika virus and microcephaly and the presence of others etiological agents. Therefore, we used a discovery-driven approach based on shotgun proteomics to acquire a comprehensive proteome of three microcephalic brains. A first strategy was to use Proteome Discoverer 2.1 and Sequest HT algorithm node to search all raw data against the entire virus entries downloaded from UniProt (April_2016). This approach robustly identified polypeptides from Bovine Viral Diarrhea Virus (BVDV). Interestingly, this strategy did not identify any Zika virus polypeptide. Afterwards, we repeated the search, now against a database containing combined virus and human entries downloaded from UniProt (April_2016). The output showed two Master Proteins (protein groups using maximum parsimony) that identified BVDV polyproteins (Q65815 and Q8B512) (Table S1). To confirm these findings, we employed a different software and algorithm, PatternLab 4.0 and Comet, respectively (9). Using the same previous virus and human database, BVDV polyproteins were also identified (Table S2). In summary, the searches identified at least 25 peptides that map to BVDB polyproteins (Table S3) and none to Zika.

These peptide sequences submitted on-line to Blastp analyses against UniProt viruses’ databank identified as first hits BVDV polyproteins with different accession numbers: Q8B512, Q8B513 and O57114 for BVDV mucosal disease virus, Q9WR85 for BVDV 2, Q9MZG9 BVDV 1 strain CP821 and BVDV 1-Osloss (Table S3). When the Blastp strategy was extended to the UniProt Human Protein database the same peptides, except one, showed 100 % matches to five different proteins: gamma-aminobutyric acid receptor-associated protein-like (P60520, GABARAP), polyubiquitin-B (J3QSA3, UBB), neural precursor cell expressed developmentally down-regulated protein 8 (Q15843, NEDD8), ubiquitin-60S ribosomal protein L40 (P62987, UBA52) and small ubiquitin-related modifier 2 (P61956, SUMO 2).

Interestingly, a unique 21-residue peptide pinpointed by the Proteome Discovery search, TLTGRTITLEVEPSDTIENVK, matched 100% of the sequence of residues 1596 to 1616 of the genome polyprotein of bovine viral diarrhea virus 1-Osloss (Q65815). Importantly, this peptide, highlighted in Table S3, showed a lower match of 95.2% to human ubiquitin-60S ribosomal protein L40 (P62987), differing from all other 24 peptides that matched 100% with both human and virus proteins. The difference is due to an exchange of R in the virus protein for a K residue in the human protein. This mutation in TLTGKTITLEVEPSDTIENVK accounts for the 100 % matches to both human ubiquitin-60S ribosomal protein L40 (P62987) and BVDV 2 polyprotein (Q9WR85). Also important is that two different PSMs for these peptides (Figure 1) reinforce the identification of peptide TLTGRTITLEVEPSDTIENVK as unique for BVDV 1-Osloss, as previously shown in the Proteome Discoverer 2.1 result. Spectra annotations confirmed the two sequences (Figure 1). The R peptide does not have 100 % match with any human protein. This made us to suspect the presence of another etiological agent, possibly a virus containing structural features of a BVDV-simile strain 1 in the three brain extracts whereas the presence of strain 2 was impossible to ascertain. Other peptides – AKIQDKEGIPPDQQR and LIFAGKQLEDGR – matched down-stream segments of the polypeptide chain of both BVDV and human ubiquitin confirming the existence and localization in this polypeptide region. BVDV is a common contaminant of bovine biological products (10, 11). To test the contamination of brain extracts by BVDV from reagents we submitted the extraction solution containing sodium deoxycholate, porcine trypsin and inhibitors to the same analytical protocol used for protein extraction, except acetone precipitation to avoid protein losses. Shotgun mass spectrometry failed to detect any peptide described in Table 1. Reports of incorporation of animal host proteins by BVDV exist since 1989 (12). Examples are the cellular protein sequences inserted in BVDV genome of segments of polyubiquitin (13), recombination events of the NS2-3 region transforming a non cytophatic to a cytophatic virus (14), ribosomal protein S27 (15), light chain 3 (LC3) of microtubule-associated proteins 1A and 1B (16), SMT3B (17), NEDD8 (18) and the J domain protein (19). Other inserts were segments of gamma aminobutyric acid (A) receptor associated protein [GABARAP] and cellular Golgi-associated ATPase enhancer (GATE-16) (20). It is not possible to ascertain whether the peptides that identified GABARAP, UBB, NEDD8, UBA52 and SUMO 2 came from human brain tissue, form BVDV genome polyprotein or both. However, it is acceptable to postulate that the published data do not invalidate our BVDV proteomics data but, on the contrary, justifies a possible scenario according to these previous works.

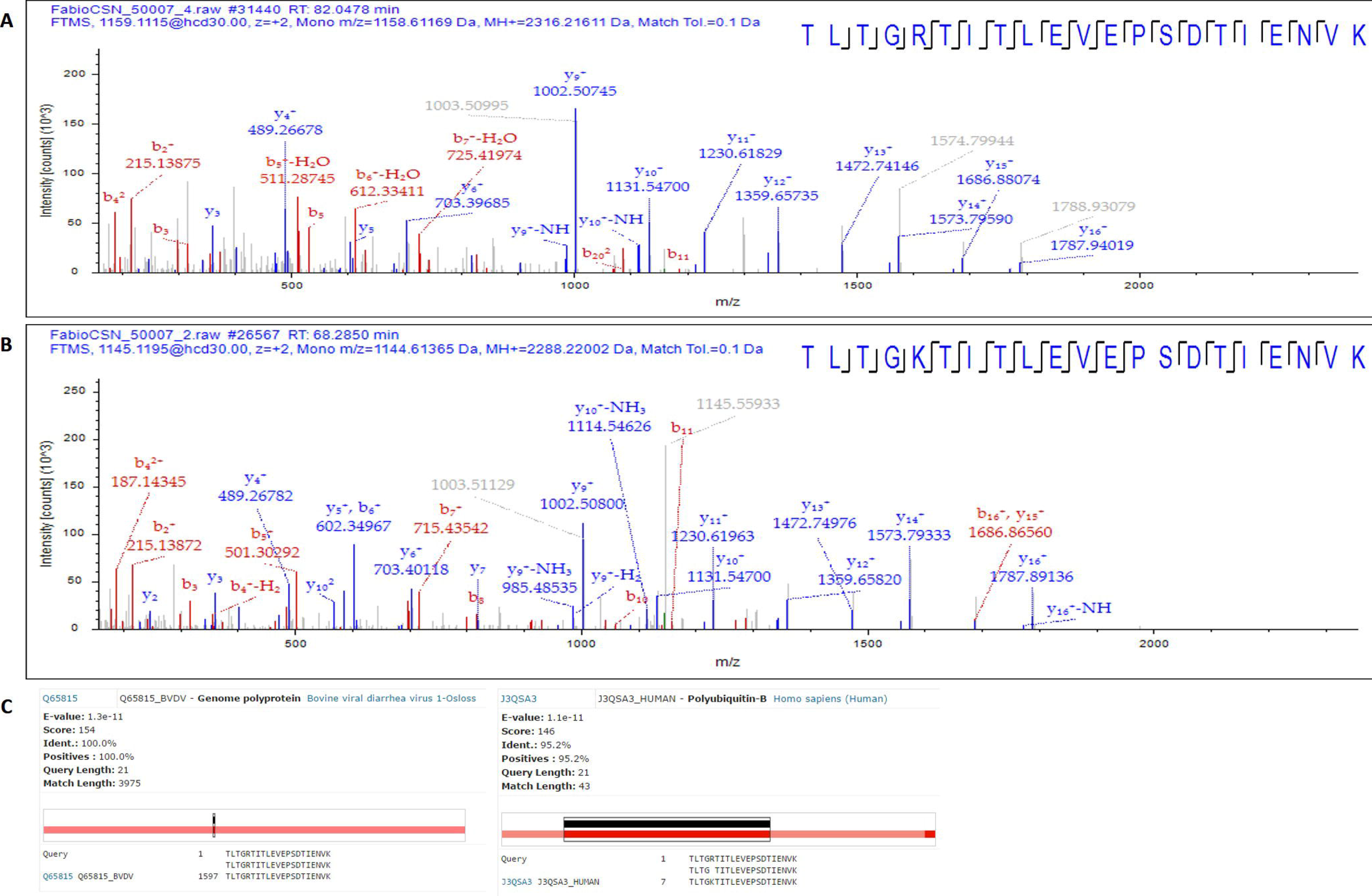

BVDV is an enveloped pestivirus in the family Flaviviridae with a single strand RNA genome of positive polarity, approximately 12.5 kb in size containing a single open reading frame flanked by two non-translating regions named 5´and 3-UTR. The single translated polypeptide chain generates 10 - 12 mature viral proteins, structural and nonstructural. Pathological lesions in animals caused by BVDV infections vary in type and degree and consequently promote wide pathogenesis including congenital defects. The virus infects the gastrointestinal tract and lymphoid tissues, the cerebellum of midgestation and placenta of aborted fetuses. Brain lesions typically consist of hydranencephaly, porencephaly, hydrocephalus and cerebellar hypoplasia as well as arthrogryposis. BVDV infection also leads to early embryonic death, abortion, fetal PI, and congenital abnormalities in cow, goats and other even ungulates (21, 22). These characteristics make a BVDV-like pestivirus a possible co-infecting agent acting together with Zika virus for the development of congenital malformations. Human fetuses with microcephaly show similar lesions (5, 7).

Reports of BVDV presence in humans are scarce. Giangaspero*et al* described in 1988 (23) the detection of specific antibodies to bovine viral diarrhea virus in high-risk animal handlers, those that had direct contact with potentially infected animals. Monoclonal-antibody-based immunoassay detected Pestivirus antigens in feces of children with gastroenteritis (24) and the virus strain Europa (NCP) identified as BVDV by radioimmunoassay was isolated from a human buffy coat from human leucocytes in Belgium (25). In the USA 40% of sera from identical twins discordant for schizophrenia tested positive for specific anti-BVDV antibodies (26). Infection during pregnancy and a possible association with some forms of cerebral White Matter Damage (WMD) in preterm neonates have been investigated (27).

Animal and herd infection by BVDV in the State of Paraíba, Brazil have been reported. Spatial hierarchical variances and age co-variances for seroprevalence to BVDV sampled from 2343 cattle from 72 properties showed seroprevalence to BVD at 22.2 % strongly clustered within herds and positively correlated between young and replacement age groups within herds. Another cross-sectional study carried out from September 2012 to January 2013 in 2443 animals sampled from three strata showed the seroprevalence of seropositive animals to be 65.5% at 95% confidence interval (CI) for herd and 39.1% (95% CI⦋=⦋33.1-45.6%), for animal, whereas the frequency of seropositive animals per herd ranged from 10 to 100% (median of 50%) with a risk factor of more than six calves aged ≤12 months of 3.72; 95% CI (28, 29).

In conclusion, our data made us to suspect that Zika virus may not be the sole etiological agent present in brains of fetuses born with microcephaly. In fact, the data raises the suspicion that another agent possessing similar structural features and properties to BVDV might induce the same anomalies to animal and human fetuses. Finally, it did not escape to our attention one working hypothesis for viruses functioning in humans: concerted actions in a synergistic way such that Zika virus breaks down physiological barriers for entrance of the BVDV-like virus.

## Contributors

ASOM did the clinical work and collected brain samples, EV, FCSN and GBD designed the experiments, EV and FCSN performed the analysis, FCSN and GBD analyzed the data. FCSN and GBD wrote and edited the manuscript. All authors reviewed the final draft.

## Acknowledgements

we thank the Rio de Janeiro Science Funding Agency for research grants E-26/111.697/2013, E-26/110.138/2013, E-26/010.001582/2014 and Young Scientist scholarship to FCSN E-26/ 202.801/2015) and the National Council for Scientific and Technological Development (grants # 477325/ 2013) as well as scholarships to GBD and EV (#306316/20153 and #141580/2015–1).

## BIBLIOGRAPHY

1. Campos GS, Bandeira AC, Sardi SI. Zika Virus Outbreak, Bahia, Brazil. Emerg Infect Dis 2015; 21: 1885–86.

2. Brazilian Ministry of Health.Ministério da Saúde investiga 3.852 casos suspeitos de microcefalia no paÍs (inportuguese). portalsaude.saude.gov.br/index.php/cidadao/principal/agencia-saude/22145. Ministerio da Saude investiga 3.852 casos suspeitos de microcefalia no pais (accessed Feb 11, 2016).

3. Oliveira Melo AS, Malinger G, Ximenes R, Szejnfeld PO, Alves Sampaio S, Bispo de Filippis AM. Zika virus intrauterine infection causes fetal brain abnormality and microcephaly: tip of the iceberg? Ultrasound ObstetGynecol 2016; 47: 6–7.

4. Mlakar J, Korva M, Tul N, Popović M, Poljšak-Prijatelj M, Mraz J, Kolenc M, Rus KR, Vipotnik TV, Vodušek FV, Vizjak A, Pižem J, Petrovec M,and Županc TA. Zika Virus Associated with Microcephaly. N Engl J Med 2016; 374: 951–958.

5. Rasmussen SA, Jamieson DJ, Honein MA, Petersen LR. Zika Virus and Birth Defects - Reviewing the Evidence for Causality. N Engl J Med 2016; 374: 1981–1987.

6. Garcez PP, Loiola EC, Madeiro da Costa R, Higa LM, Trindade P, Delvecchio R, Nascimento JR, Brindeiro R, Tanuri A, Rehen SK. Zika vÍrus impairs growth in human neurospheres and brain organoids. Science 2016; 352: 816–818.

7. Besnard M, Lastère S, Teissier A, Cao-Lormeau,VMC and Musso D. Evidence of perinatal transmission of Zika virus, French Polynesia, December 2013 and February 2014. Euro Surveill 2014; 19:8–11.

8. Calvet G, Aguiar RS, Melo ASO, Sampaio SA, de Filippis I, Fabri A, Araujo ESM, Sequeira PC, Mendonça MCL, Oliveira L, Tschoeke DA, Schrago CG, Thompson FI, Brasil P, Santos FB, Nogueira RMR, Tanuri A, de Filippis AMB. Detection and sequencing of Zika vÍrus from amniotic fluid fetuses with microcephaly in Brazil: a case study. Lancet Infect Dis 2016; 16: 653–60.

9. Carvalho PC, Lima DB, Leprevost VF, Santos MDM, Fischer JSG, Aquino PF, Moresco JJ, Yates III JR, Barbosa VC. Integrated analysis of shotgun proteomic data with PatternLab for proteomics 4.0. Nature Protocols 2016; 11: 102–117.

10. Studer E, Bertoni G and Candrian U. Detection and Characterization of Pestivirus contaminations in Human Live Vaccines. Biologicals 2002; 30: 289–296.

11. Giangaspero M. Pestivirus Species Potential Adventitious Contaminants of Biological Products. Trop Med Surg 2013; 1: 153. doi:10.4172/2329-9088.1000153.

12. Meyers G, RÜmenapf T and Thiel H-J. Ubiquitin in a Togavirus. Nature 1989; 341: 491.

13. Meyers G, Tautz N, Dubovi EJ and Thiel H-J. Viral Cytopathogenicity Correlated with Integration of Ubiquitin-Coding Sequences. Virology 1991; 180: 602–616.

14. Deng R and Brock KV. Molecular Cloning and Nucleotide Sequence of a Pestivirus Genome, Noncytopathic Bovine Viral Diarrhea Virus Strain SD-l. Virology 1992; 191: 867–879.

15. Becher P, Orlich, M and Thiel H-J. Ribosomal S27a Coding Sequences Upstream of Ubiquitin Coding Sequences in the Genome of a Pestivirus. J, Virology 1998; 72: 8697–8704.

16. Meyers G, Stoll D and Gunn M. Insertion of a Sequence Encoding Light Chain 3 of Microtubule-Associated Proteins 1A and 1B in a Pestivirus Genome: Connection with Virus Cytopathogenicity and Induction of Lethal Disease in Cattle. J Virology 1998; 4139–4148.

17. Qi F, Ridpath JF and Berry ES. Insertion of a bovine SMT3B gene in NS4B and duplication of NS3 in a bovine viral diarrhea virus genome correlate with the cytopathogenicity of the virus. Virus Res 1998; 57: 1–9.

18. Baroth M, Orlich M, Thiel, H-J and Becher P. Insertion of Cellular NEDD8 Coding Sequences in a Pestivirus. Virology 2000: 456–466.

19. Müller A, Rinck G, Thiel H-J, and Tautz N.. Cell-Derived Sequences in the N-Terminal Region of the Polyprotein of a Cytopathogenic Pestivirus. J Virology 2003; 77: 10663–10669.

20. Becher P, Thiel H-J, Collins M, Brownlie J and Orlich M. Cellular Sequences in Pestivirus Genomes Encoding Gamma-Aminobutyric Acid (A) Receptor-Associated Protein and Golgi-Associated ATPase Enhancer of 16 Kilodaltons. Journal of Virology 2002; 76:13069–13076.

21. Brodersen BW. Bovine viral diarrhea virus infections. Manifestations of Infection and Recent Advances in Understanding Pathogenesis and Control. Veterinary Pathology 2014; 51:453-4641-12.

22. Webb BT, Norrdin RW, Smirnova NP, Campen V, Weiner CM, Antoniazzi AQ, Bielefeldt-Ohman H,and Hansen TR. Bovine Viral Diarrhea Virus Cyclically Impairs Long Bone Trabecular Modeling in Experimental Persistently Infected Fetuses. Veterinary Pathology 2012; 49: 930–940.

23. Giangaspero M, Wellemans G, Vanopdenbosch E, Belloli A and Verhulst A. Bovine viral diarhhoea. The Lancet 1988; 332: 110.

24. Yolken R, Dubovi E, Leister F, Reid R, Almeido-Hill JK and Santrosham M. Infatile gastroenteritis associated with excretion of pestivirus antigens. The Lancet 1989; 1(8637): 517–520.

25. Giangaspero M, Harasawa R, Verhulst A. Genotypic analysis of the 5’-untranslated region of a pestivirus strain isolated from human leucocytes. MicrobiolImmunol 1997; 41: 829–834.

26. Yolken RH, Petric M, Collett M, Fuller TE. Pestivirus infection in identical twins discordant for schizophrenia. Proceedings of the IXth Congress of Virology, Glasgow, England 148, 1993.

27. Rennie J, Peebles D. Pestivirus as a cause of white matter damage down but not out. Dev Med Child Neurol 2006; 48: 243.

28. Thompson JA, de Miranda HLR, Gonçalves VS, Leite RC, Bandeira DA, Hermann GP, Moreira EC, Prado PE, Lobato Zi, de Brito CP and Lage AP. Spatial hierarchical variances and age covariances for seroprevalence to Leptospira interrogans serovar hardjo, BoHV-1 and BDVD for cattle in the State of ParaÍba, Brazil. Prev. Vet. Med. 2006; 76:290–301

29. Fernandes LG, Nogueira AH, De Stefano E, Pituco EM, Riberiro CP, Alves CJ, Oliveira TS, Clementino IJ and de Azevedo SS. Herd-level prevalence and risk factors for bovine viral diarrhea vÍrus infection in cattle in the State of Paraíba, Northeastern Brazil. Trop Anim Health Prod 2016; 48: 157–165

